# The role of Miocene-Pleistocene environmental change in diversification of the genus *Hypoclinemus* Chabanaud, 1928

**DOI:** 10.1101/2023.12.07.570667

**Authors:** Alan Erik S. Rodrigues, Jonathan Stuart Ready, Lucas Gabriel Pereira da Silva, Kamila de Fátima Silva, Derlan José Ferreira Silva, Mark H. Sabaj, Marcelo C. Andrade, Santelmo Vasconcelos, João Bráullio L. Sales

## Abstract

The Neotropical region stands out as one of the most taxonomically diverse areas on the planet, garnering significant attention in the context of marine incursions and their role in shaping this diversity. Among marine-derived taxa, pleuronectiform fishes exhibit distinctive morphological characteristics that have attracted significant scientific interest. However, the biogeography of *Hypoclinemus mentalis*, initially described as *Solea mentalis* and subsequently assigned to the genus *Achirus*, was eventually reclassified into its current monotypic genus due to its limited distribution in freshwater environments, in contrast to the species of *Achirus*. The broad distribution of a single species across multiple South American river basins positions *H. mentalis* as an ideal candidate for biogeographic studies within South America, with an emphasis on the detection of cryptic lineages associated with major drainage basins. In our study, we employed mitochondrial and nuclear markers to investigate the potential existence of such lineages within the broader context of a molecular phylogeny that encompasses all valid genera in the flatfish family Achiridae. Our findings reveal that *Hypoclinemus* comprises seven operational taxonomic units (OTUs), as deduced from specimens collected across the majority of its documented range. Furthermore, our phylogeographic analyses support the hypothesis that colonization of freshwater habitats occurred through connections between the Caribbean Sea and Lake Pebas approximately 21.28 million years ago. Moreover, we observed that differentiation of lineages within the *Hypoclinemus* genus was significantly influenced by pronounced sea level fluctuations during the Plio-Pleistocene epoch, underscoring the impact of glaciations and interglacial periods on the biogeographic patterns.

## Introduction

The neotropical region is one of the world’s most taxonomically diverse areas especially considering its ichthyofauna that includes more than 6200 known species [1] and estimates of 8000 to 9000 total species [2, 3]. Neotropical fishes include several groups that have adapted to freshwater environments from marine origins such as rays (Potamotrygoninae, [4], pufferfish (Tetraodontidae, [5]), needlefish (Belonidae, [6]), drums and anchovies (Sciaenidae and Engraulidae, respectively, [7, 8]) and flatfishes, family Achiridae [9]. The adaptation of these taxa to freshwater environments is generally attributed to marine incursions [4, 6, 8], especially those associated with the Pebas system, a mega-wetland which formed during the latest Oligocene to Early Miocene (∼24 to 16 mya) and reached its maximum extent during the Middle to early Late Miocene (∼16 to 11.3 mya) [10, 11]. Alternatively, some marine groups may have adapted to freshwater environments via estuarine adaptation, such as silversides in the Old (Atherinidae) and New (Atherinopsidae) Worlds [12,13].

For most of the Miocene, the Pebas system experienced a connection to the Caribbean Sea, which may have offered an ideal transitional environment for adaptations and colonization of proto-marine lineages in freshwater environments, especially in the region that now comprises the Amazon [14, 15]. Subsequently, the diversification of taxa of marine origin depends on speciation processes common to continental freshwaters. In the case of the Amazon, periods of glaciation during the Pleistocene and tectonic processes associated with the uplift of the Andes resulted in changes in abiotic conditions and drainage connections in the region [4, 16, 17, 18]. However, for other families that have both marine and freshwater species (e.g., Achiridae), there is little information about the influence of these past processes of speciation [19].

The flatfish family Achiridae is a member of the superfamily Soleoidea, suborder Pleuronectoidei, Order Pleuronectiformes [9, 20, 21]. Although some molecular studies have questioned the monophyly and ordinal status of Pleuronectiformes (see references in Atta et al. [22]. Achiridae is composed of six genera: *Achirus* Lacépède, 1802, *Apionichthys* Kaup, 1858, *Gymnachirus* Kaup, 1858, *Catathyridium* Chabanaud, 1928, *Trinectes* Rafinesque, 1832, and *Hypoclinemus* Chabanaud, 1928. Some achirids are exclusive to marine environments (*Gymnachirus*), while others are entirely restricted to freshwater (*Hypoclinemus*). Other achirids are predominantly freshwater (*Apionichthys* and *Catathyridium*) and some (*Achirus* and *Trinectes*) inhabit multiple environments across a range of salinities [9]. The phylogenetic relationships of Achiridae have been debated in recent years, with some studies placing it near Citharidae [19, 23] or Rhombosoleidae [24, 25]. The placement of Achiridae within Soleoidea is uncertain due to disagreement over the composition of this superfamily [22]. Betancur-R et al. [24] and Harrington et al. [26] support Achiridae as the first lineage to diverge within a clade that also includes Achiropsettidae, Paralichthodidae, and two clades of Rhombosoleidae (I and II); that larger clade is sister to one composed of the remaining members of Soleoidea (Cynoglossidae, Poecilopsettidae, Samaridae, Soleidae). Atta et al. [22], however, transferred Achiropsettidae, Paralichthodidae, and Rhombosoleidae to a separate superfamily (Pleuronectoidea), and supported Achiridae as sister to a clade composed of Cynoglossidae, Poecilopsettidae, Samaridae, and Soleidae within a restricted Soleoidea. Recently, Bitencourt et al. [27] showed that Achiridae is indeed a monophyletic family that arose during the Oligocene-Miocene transition.

Bitencourt et al. [27] also point out that some genera may harbor cryptic lineages and that *Hypoclinemus* may have a closer relationship with the samples of *Achirus achirus* (Linnaeus, 1758) used in their study. Chabanaud [28] proposed *Hypoclinemus* for two species, *H. paraguayensis* which he described from the Paraguay river, and the nominal *Solea mentalis* described by Günther (1862) from the Capim river near Belém, Brazil. Ramos [29] synonymized the former species with *Catathyridium lorentzii* (Weyenbergh, 1877). Described as *Achirus hasemani* (Steindachner, 1915) from Rio Branco, Brazil, which was also synonymized with *H. mentalis* [28]. Thus, *Hypoclinemus* is monotypic with only *H. mentalis* considered valid. *Hypoclinemus* is distinguished by having teeth on both rami of the dentary vs. restricted to the left ramus (blind side) in other achirids [21, 30]. This characteristic is considered the most plesiomorphic among achirids, providing osteological support for *H. mentalis* as the sister species to the rest of the group. *Hypoclinemus mentalis* is widely distributed throughout Greater Amazonia, which includes the Amazon, Essequibo, and Orinoco basins [31]. Its large distribution raises questions as to whether *Hypoclinemus* is in fact monotypic, since other widespread neotropical fishes have been recently split into multiple species (e.g., De Santana et al. [32]; Garavello et al. [33]) or shown to include hidden diversity represented by multiple species-level lineages (e.g., Mendes et al. [34]).

Given the complex topology of Greater Amazonia (sensu Van der Sleen and Albert, [31]), *Hypoclinemus* is also a good candidate to investigate how speciation processes may affect the evolutionary history of Neotropical fish species, which in many cases include the divergence of cryptic lineages [18, 35, 36, 37]. In freshwater environments, fishes found in smaller streams and headwaters tend to be more strongly fragmented and isolated, even on small spatial scales, providing ample opportunity for speciation [18, 38], while fishes common to major rivers with accumulated sedimentary substrates tend to speciate based on large-scale vicariance events between basins and ecological differences across basins [32]. The geographic range of *H. mentalis* spans major basins (Amazon, Orinoco, Essequibo) that present spatial and environmental heterogeneity, factors that can influence speciation [39]. Therefore, this study aims to test the monophyly of the genus *Hypoclinemus,* exploring whether *H. mentalis* is composed of a single evolutionary lineage across its distribution in the Neotropics, or whether multiple lineages exist that can create a biogeographic profile for understanding the history of the region.

## Material And Methods

### Tissue Sampling

Muscle tissue samples were obtained from specimens of *Hypoclinemus* (n=27) and *Apionichthys* (3) distributed among seven ecoregions (Fig. 1, S1 Table).

**Fig 1.**
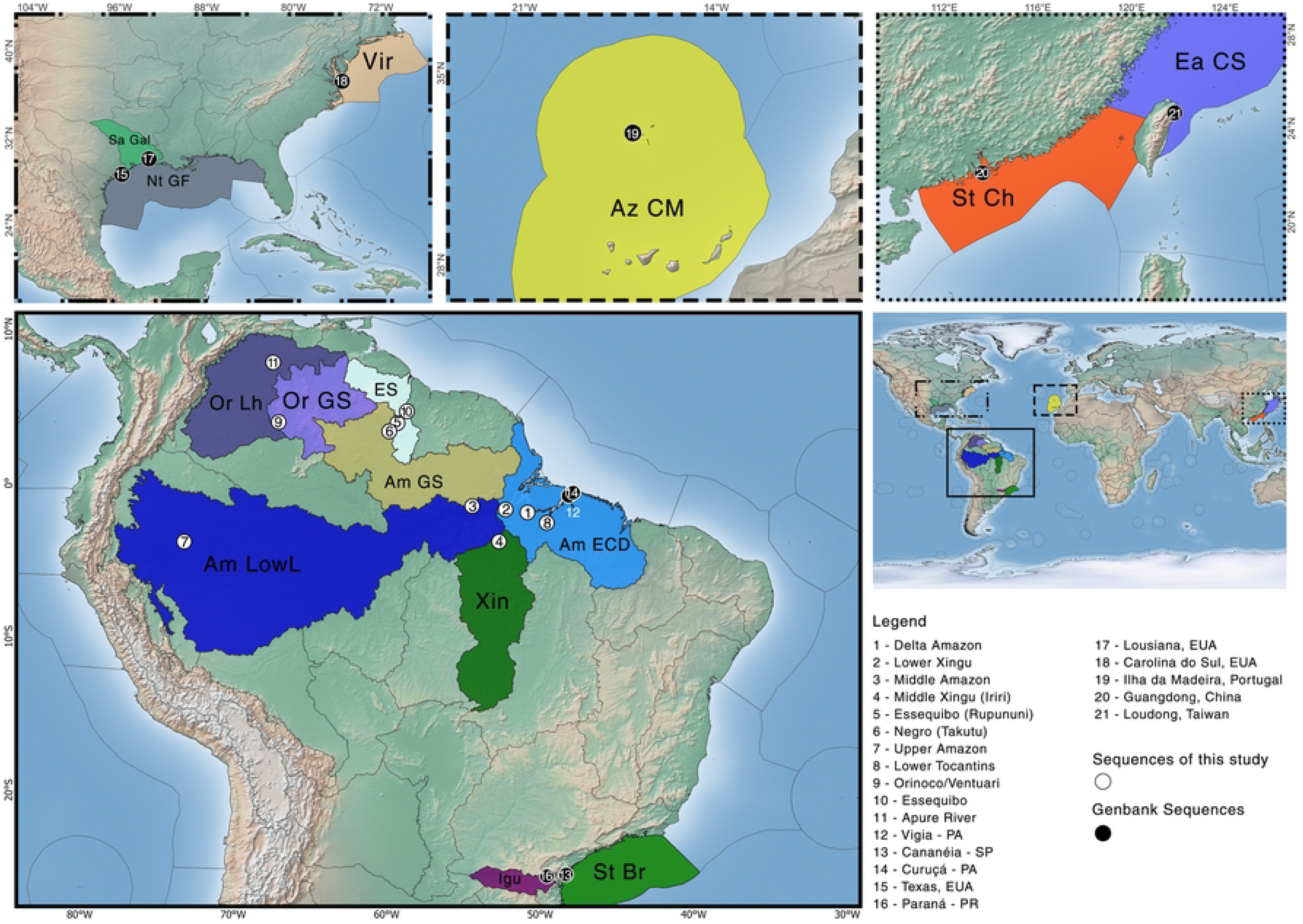
Map showing the distribution of newly sequenced samples and previously existing GenBank data analyzed in the present study. The ecoregions associated to each sample are based on Spalding et al. [52] and Abel et al. [53], using the following abbreviated names: Am ECD = Amazonas Estuary and Coastal Drainages; Am GS = Amazonas Guiana Shield; Am LowL = Amazonas Lowlands; ES = Essequibo; Or GS = Orinoco Guiana Shield; Or Lh = Orinoco, Llanos; Xin = Xingu; Az CM = Azores Canaries Madeira; Ea CS = East China Sea; Igu = Iguassu; Nt GF = Northern Gulf of Mexico; Sa Gal = Sabine-Galveston; St Br = Southeastern Brazil; St Ch = Southern China; Vir = Virginia. The numbers indicate the locations of the samples. The different colors indicate the ecoregions in RASP 4.

Two specimens of *Hypoclinemus mentalis* were collected in the sandy beach region of Portel - PA (Delta, Amazon - Ecoregion of the Amazon estuary and coastal drainages) and one in Mocajuba - PA (Lower Tocantins - Ecoregion Amazon estuary and coastal drainages), in addition to three individuals from Alenquer – PA (Middle Amazon – Guiana Shield Ecoregion of the Amazon). Specimen vouchers were deposited in the collection of the CEABIO (Centre for Advanced Biodiversity Studies - UFPA). Samples were collected under the SISBIO license number 53022-2 from the “Instituto Chico Mendes de Conservação da Biodiversidade” and for “Sistema Nacional de Gestão ao Patrimônio Genético e do Conhecimento Tradicional Associado” (SisGen) license number AB 2492E. All work performed in accordance with ethical approval by the Federal University of Pará Committee for the Ethical Use of Animals (CEUA 68-2015). From each individual, a sample of muscle tissue was taken, stored in absolute ethanol, and kept at −4°C until DNA extraction. Additional samples were analyzed from specimens previously collected in Guyana, Peru, Venezuela, and Brazil (Xingu Basin) available in existing collections (S1).

### DNA Extraction, PCR and Sequencing

Total DNA was extracted from muscle tissue preserved in ethanol following a modified CTAB protocol [40] using only chloroform and isoamyl alcohol (CIAA) in a 1:1 proportion. For the present study, two mitochondrial markers, Cytochrome C Oxidase subunit I (*cox1*) and rrln (the mitochondrial large ribosomal subunit, also known as 16S rRNA), and one nuclear gene (Rhodopsin - *RHO*) were used (Table 1).

**Table 1.**
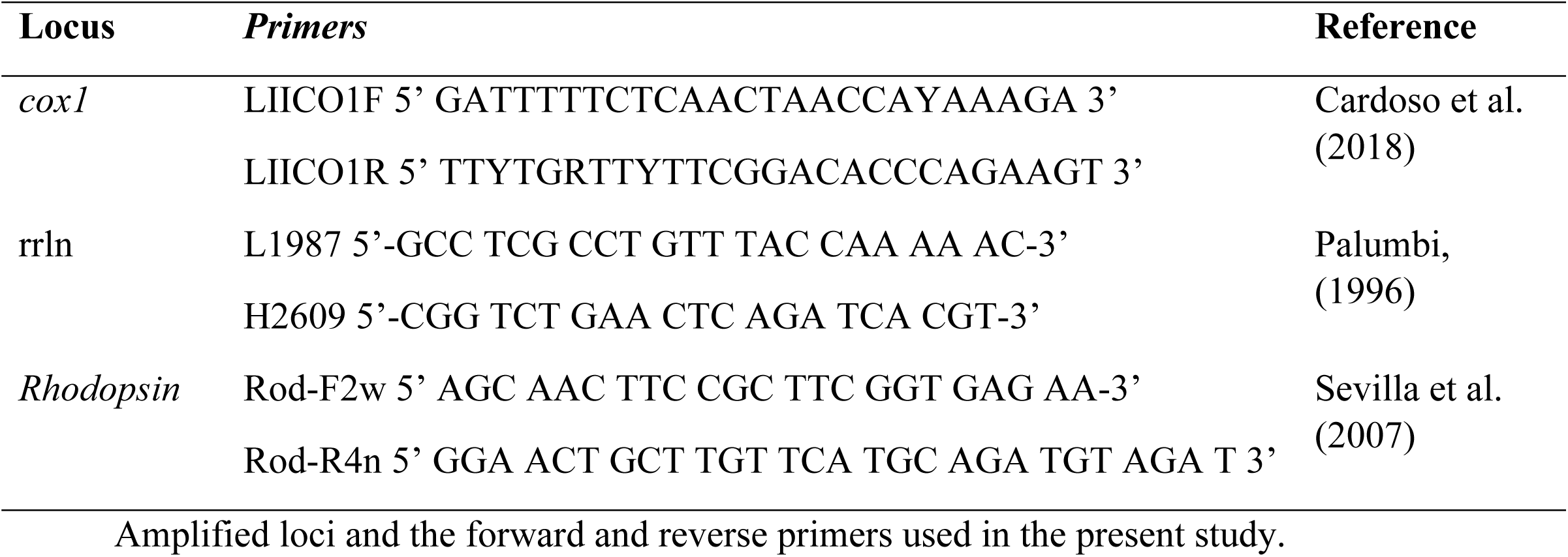
Primers used for amplifying targeted gene regions.

Each PCR reaction had a final volume of 25 μL, containing 2 mM dNTPs, Buffer (5x), 2.4 mM MgCl2, 4 μL of template DNA, 1 μL of each primer, 0.2 μL of Taq polymerase, and 10.4 μL of ultrapure (UP) water. The PCR conditions for the three markers were similar, changing only the annealing temperatures of each primer pair: 95 °C for 5 min for initial denaturation; 30 cycles of 94 °C for 40 s, 55 °C (rrln), 58 °C (*cox1*), or 63 °C (*RHO*) for 40 s, and 72 °C for 30 s; and a final extension step at 72 °C for 7 min.

PCR purifications were performed using 65% isopropanol; after two washes with 70% ethanol, all samples were resuspended in 20 μL of UP water and stored in a −20° freezer. The sequencing reaction was carried out by the terminal dideoxynucleotide method [41] using reagents from the BigDye Terminator v3.1 Cycle Sequencing kit (Applied Biosystems/Life Technologies). DNA sequences were generated in an automatic sequencer model ABI 3730 96 - capillary DNA Analyzer, from Applied Biosystems.

### Alignment and Molecular Procedures

The present study also incorporated public sequences from GenBank [42] for the three amplified markers for all six achirid genera (*Achirus*, *Apionichthys*, *Gymnachirus*, *Catathyridium*, *Trinectes*, and *Hypoclinemus*) as well as the outgroup genera *Bothus* Rafinesque, 1810, and *Symphurus* Rafinesque, 1810 (S1 Table). For our study, we used three different datasets. First, the concatenated dataset for all the three markers (rrln+*cox1*+*RHO* - dataset 1). This dataset only lacked the *cox1* sequences from Bitencourt et al. [27] that were not available in Genbank. With dataset 1, we generated phylogenies (Maximum likelihood-ML and species tree) and estimated the Time to Most Common Ancestor (TMRCA) and Reconstruction of Ancestral States (RASP 4). We also generated phylogenies using only the *cox1* data (ML and Bayesian trees - dataset 2, S2 Figure 2A) and using a concatenation of the rrln and *RHO* sequences (ML and species trees - dataset 3, S2 Figure B).

**Fig 2.**
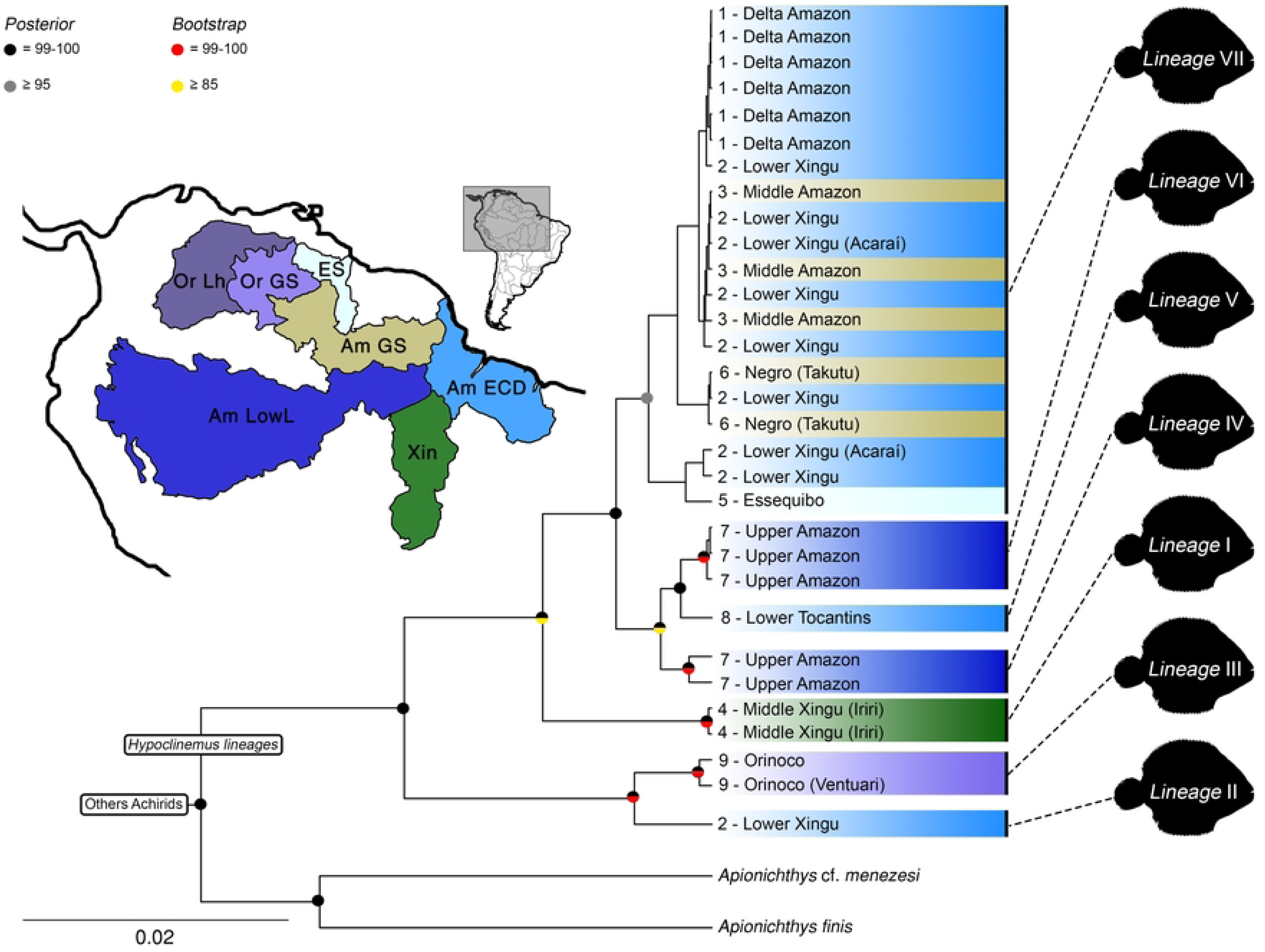
Species tree generated using *BEAST. Only support values above 0.95 for Bayesian Inference are shown, for maximum likelihood, only supports above 85% are indicated. Color of bars represent ecoregion associations inferred in RASP 4. Tip nodes are numbered according to location numbers in Fig. 1.

For all phylogenetic analyses performed in this study, we used concatenated datasets; however, evolutionary models were independently estimated for each partition using ModelFinder 2 [43], implementing the AIC criterion to determine the best evolutionary model for both Maximum Likelihood (ML) and Bayesian Inference (BI) analyses. The models GTR+I+G (rrln), HKY+I+G (*cox1*), and HKY+I (*RHO*) were selected for both ML and BI analyses for dataset 1. For dataset 2 (*cox1*), TIM2+I+G (ML) and HKY (BI) were selected. Finally, for dataset 3 (rrln+*RHO*), TIM2+I+G (ML and BI) and HKY+I+G (ML and BI) were selected, respectively.

Initially, an ML tree was generated in IQ-TREE 2 [44] using the 1000 ultrafast bootstrap replicates [45]. A species tree was then produced using *BEAST [46] which is part of the BEAST v.1.7.4 package [47]. In BEAUTi, the evolutionary models were “unlinked” to use the previously estimated models for each separate partition. Only species with sequence data for more than two markers were analyzed. For this, we used an uncorrelated relaxed clock with a lognormal distribution prior on rates and a Yule speciation prior [46, 48]. MCMC (Markov Chain Monte Carlo) samples were estimated through four simultaneous runs containing four chains (one cold and three warm) with 50 million generations. All posterior probabilities were defined using the 80% consensus rule, with samples taken every 1000 generations, and 10% of the initial trees discarded as burn-in. Log-likelihood files generated in each run were viewed in Tracer v.1.4 [47], where only runs with ESS values equal to or greater than 200 were considered. The consensus tree was then generated using TreeAnnotator v.1.4 [47].

### Divergence Time Estimates and Biogeographic Reconstruction

The time to The Most Recent Common Ancestor (TMRCA) was estimated with BEAST 2 [49], using the three previous independent partitions. The TN93+G model was used with base frequencies determined empirically. For tree prior parameters, the Yule model was selected, using the uncorrelated relaxed clock and a lognormal distribution for rates without enforcing monophyly for any genera [48]. We used two dating priors in the BEAST2 analysis, both of which were estimated in Shi et al. [23], based on fossil calibrations from Near et al [50]. The first one is the separation between *Samaris cristatus* Gray, 1831, and *Poecilopsetta beanii* (Goode, 1881) at 52.9 million years ago (mya). We assigned a log-normal distribution prior and set this prior to have an offset of 51.0, standard deviation (S) of 0.5, and mean (M) of 17.78. For the second calibration, we used the separation between *Trinectes* and *Achirus* at 18 mya, where we assigned a log-normal distributed prior and set this prior to have an offset of 15.0, standard deviation (S) of 0.5, and mean (M) of 16.7. The MCMC method was used to infer the divergence times with four independent runs with 120 million generations through four simultaneous runs containing four chains (one cold and three heated) with sampling performed every 1000 generations. Only runs with ESS values equal to or greater than 200 for all marginal parameters were used. Log-likelihood files generated in each run were viewed in Tracer v.1.4 [47], where only runs with ESS values equal to or greater than 200 were considered. The consensus tree was then generated using the TreeAnnotator v. 1.4 [47].

RASP 4 [51] was used to reconstruct the ancestral biogeographic area, using the Bayesian Binary Method (BBM). The TMRCA tree generated in BEAST 2 was used as the guide tree. We used the ecoregion level areas defined for marine [52] and freshwater [53] species. The BBM analyses were run for eight million cycles, using ten chains, with sampling every 100 cycles. The temperature was set to 0.1, and a fixed JC model was used. The maximum number of areas for all nodes was set at four. Subsequently, the information of each node was plotted on pie charts.

## Results

### 3.1 Phylogenetic Reconstructions

A total of 30 individuals were sequenced for each of the three genes (rrln, *cox1*, and *RHO*) in the present study. All GenBank sequences utilized for each dataset are listed in S1 Table. Together these resulted in a total of 187 individuals for analyses.

Based on the phylogenetic reconstructions simulated here (based on two mitochondrial regions and one nuclear gene) *Hypoclinemus* was recovered as a monophyletic genus, and as sister taxon to *Apionichthys* with a high posterior probability (Fig. 2).

We also recovered seven distinct lineages within *Hypoclinemus*. Lineage I is from the Lower Xingu (Amazonas Estuary and Coastal Drainages ecoregion). Lineage II is composed of two specimens from the Orinoco basin (Orinoco Guiana Shield ecoregion) with high support values (Fig. 2). Lineage III is composed of two specimens from the Iriri River (Xingu ecoregion). Lineage IV is composed of two specimens from the Upper Amazon (Iquitos, Amazonas Lowlands ecoregion). Lineage V is represented by a single specimen from the Lower Tocantins (Amazonas Estuary and Coastal Drainages ecoregion). Lineage VI is composed of six specimens, including three from the Upper Amazon (Iquitos and Amazonas) and three from the middle Amazon (Amazonas) (Amazonas Lowlands ecoregion). Lineage VII is composed of a large clade containing the rest of the specimens utilized in the present study from many ecoregions including Essequibo, Lower Xingu, Middle Amazon and Delta Amazon. (Figure 2). Some differences in the relationships between, and support for, lineages can be found when analyzing the Dataset 2 (*cox1*) or Dataset 3 (rrnl+RHO), but are generally concordant (S2 Figure A and B). Using only Dataset 2 weakens support for most recent nodes, resulting in poor support for monophyly of the clade *Apionichthys*+*Hypoclinemus*, since *A.* cf. *menezesi* was a recovered sister to *Hypoclinemus* lineage I previously estimated here (S2 Figure A). But we also recovered significant support for four of the lineages within *H. mentalis*. A similar trend occurs when using Dataset 3 recovering also four lineages of *H. mentalis,* and for this dataset *Hypoclinemus* was recovered as a monophyletic clade (S2 Figure B).

#### Analysis of Time to Most Recent Common Ancestor and Reconstruction of Ancestral States

Three ecoregions predominated the analyses of speciation events in *Hypoclinemus*: Amazonas Lowlands, Xingu, and Amazonas Estuary and Coastal Drainages, where the latter concentrates most of the cases (Fig. 3, Table 2).

**Fig 3.**
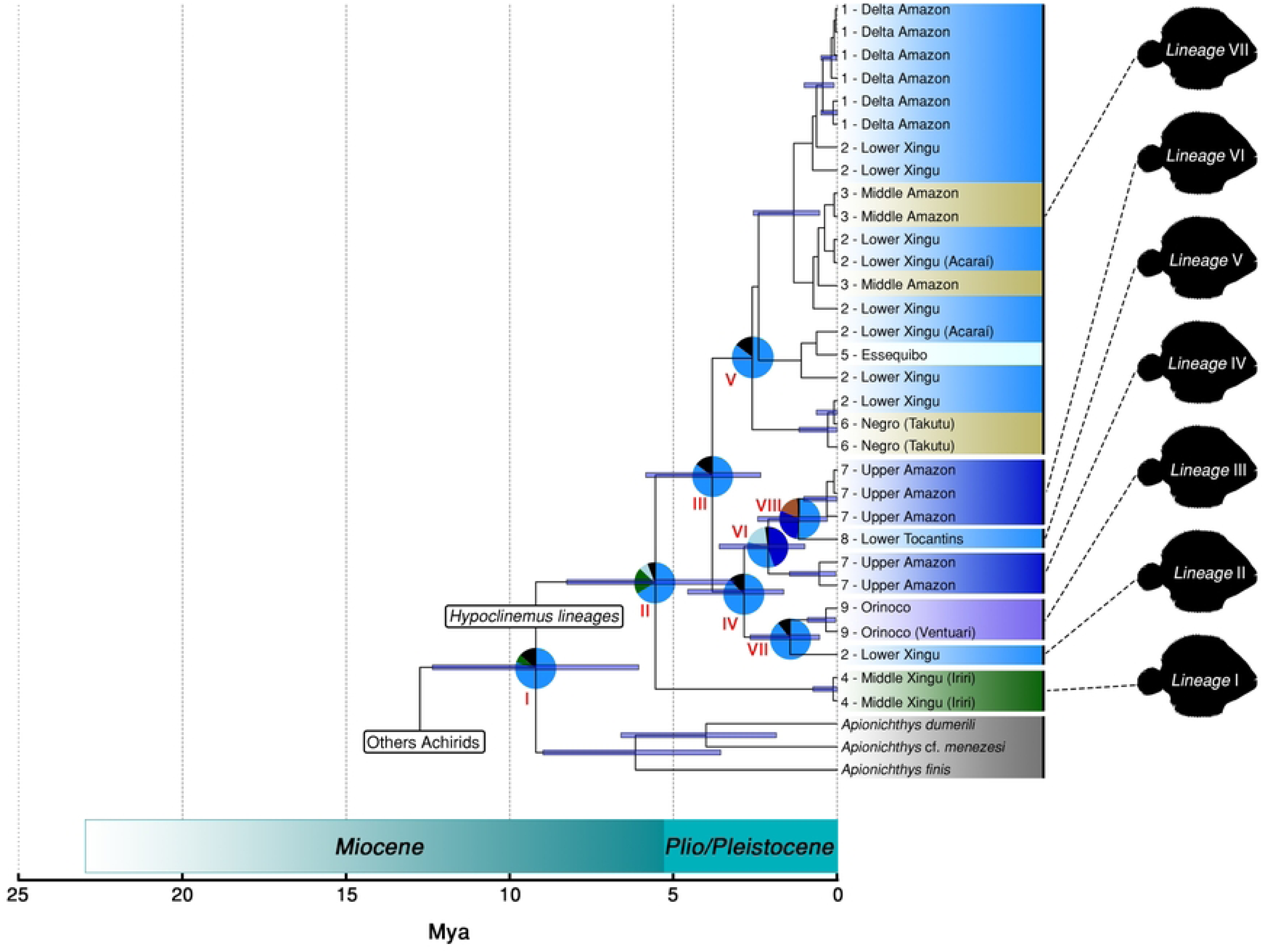
Historical biogeographical reconstruction. Inferred by BBM method in RASP 4 [51] combined with Time of Most Recent Common Ancestor tree (TMRCA) generated in BEAST 2 utilizing the log-normal relaxed clock. Blue horizontal bars on the nodes correspond to 95% High Posterior Density (HPD). Numbers indicate the split between the nodes as presented in Table 2. The different color ellipses combine the ecoregion’s description from Spalding et al. [52] and Abel et al. [53]. Lineages are numbered I-VII on the right side of the phylogeny. Tip nodes are numbered according to location numbers in Fig. 1.

**Table 2:**
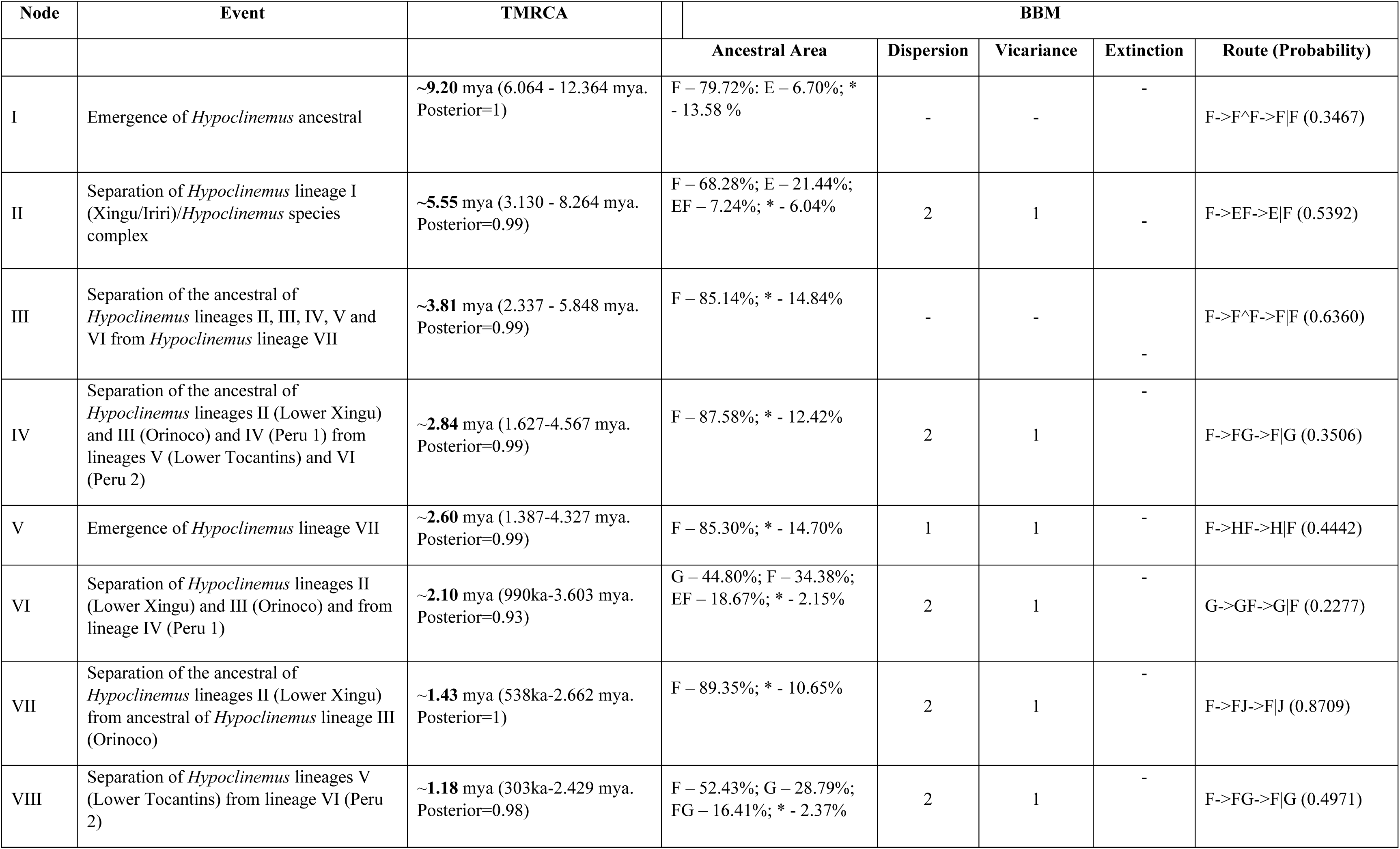
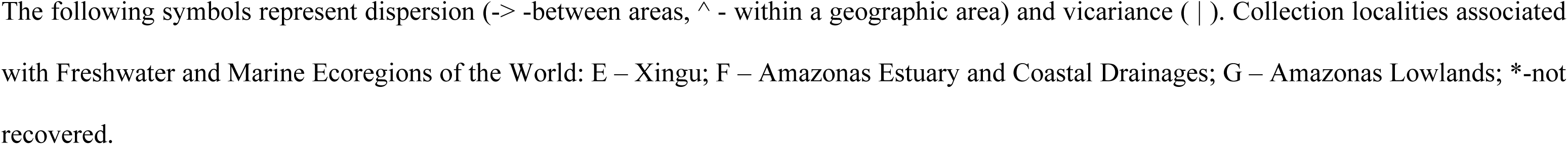
Summary of RASP 4 (BBM) ancestral area reconstructions.

Our TMRCA shows that *Hypoclinemus* split from its common ancestor with *Apionichthys* in the Miocene (∼9.20 mya), most probably in the Amazon Estuary and Coastal Drainages (72.72%) and Xingu (6.70%) ecoregions (Fig. 3, Table 2, Node I). The end of the Miocene/beginning of the Pleistocene marked the rise of the first distinct lineage (Lineage I) inside *Hypoclinemus*, diverging from the common ancestor of all *Hypoclinemus* at ∼5.55 mya, probably in the same ecoregions (Amazon Estuary and Coastal Drainages and Xingu) but resulting from … two events of dispersion and one vicariance (Fig. 3, Table 2, Node II). The subsequent divergence of Lineage VII from its shared common ancestor (the ancestor of lineages II, III, IV, V, and VI) is estimated to have taken place at ∼3.81 mya. The common ancestor of lineages II and III is then estimated to have divergence from its shared common ancestor (the common ancestor of lineages IV, V, and VI) at ∼2.84 mya (nodes III and IV, respectively, in Fig.3, Table 2).

In a short interval of time, the TMRCA and RASP analyses (Fig. 3, Table 2) estimated divergence within lineage VII at ∼2.60 mya (node V) and the separation of lineage IV from its common ancestor with the common ancestor of lineages V and VI at ∼2.10 mya (node VI). Finally, in the Pliocene, these analyses estimated the divergence between lineage I from Lower Xingu and lineage II from Orinoco at ∼1.43 mya and the divergence of lineage V from the Tocantins and VI from the Upper Amazon at ∼1.18 mya. (Nodes VII and VIII, Fig. 3, Table 2).

## Discussion

### Phylogenetic patterns within *Hypoclinemus* Chabanaud, 1928

Our results shed new light on hidden diversity inside the monotypic genus *Hypoclinemus*. However, the results generated here are different from Bitencourt et al. [27], most likely because of the distinct composition of our datasets in comparison with those of Bitencourt et al. [27], primarily because sequences of *cox1* from their work were unavailable. Additionally, our study employed a smaller number of molecular markers, and we used different sequences and samples compared to those utilized by those authors.

The presence of *A. achirus* within *Hypoclinemus* (rendering it paraphyletic) described in Bitencourt et al. [27] was not observed in any of the phylogenetic reconstructions generated in the present study. Instead, *A. achirus* was only recovered as a sister lineage to the *Hypoclinemus* species complex in reduced datasets and this relationship was not always strongly supported. This was observed in the topologies derived from dataset 2 (only *cox1* sequences - S2 Figure A) and dataset 3 (only rrln + *RHO* sequences - S2 Figure B). However, with dataset 1 (the most complete dataset formed by the three markers) and in our TMRCA analysis (dataset 1), *Apionichthys* was consistently recovered as the sister genus. It is worth noting that Byrne et al. [19] conducted a phylogenetic study with nine molecular markers (four mitochondrial and five nuclear) within the order Pleuronectiformes, including representatives of 13 of the 14 families. Their results, similar to ours, confirmed the validity of the genus *Hypoclinemus* at the genus level. However, there were differences in the placement of *Hypoclinemus* as a sister of *Achirus*, as opposed to our findings where the sister genus was *Apionichthys*.

Ramos [21] diagnosed *Hypoclinemus* based on the presence of teeth on both rami of the dentary, as opposed to them being restricted to the left ramus (blind side) as in other achirids. This led Ramos [21] to suggest that *Hypoclinemus* could be the earliest lineage, possibly originating in freshwater. However, our results present a contrasting scenario. *Hypoclinemus*, according to all phylogenetic reconstructions in our study, does form a monophyletic group, but it is not monotypic. Our findings indicate that it is sister to the genus *Apionichthys.* We also discovered the existence of at least seven lineages (putative cryptic species) within *Hypoclinemus*. To further clarify and confirm the presence of these species-level groups within the genus *Hypoclinemus*, additional taxonomic and systematic investigations are required.

### Historical biogeography of the genus *Hypoclinemus*

Several authors have discussed the origin and diversification of freshwater fish lineages with recent evolutionary ties to marine species. Among the most used hypotheses is marine invasion through river mouths [54]. However, according to many authors, marine incursions on the South American continent began in the Cretaceous period, but it was during the Miocene that the greatest number of neotropical freshwater taxa diverged from their marine relatives [4, 7, 8]. During the transition between Oligocene to Miocene, the proto-Amazon was draining into Caribbean region which promotes several marine intrusions associated with Pebas system generating colonization in western Amazon region, which in turn could promote biotic interchanges of faunas between marine and freshwater environments [4, 11].

The formation of the Pebas mega-wetlands, stretching from the Caribbean to southern South America [11, 14], played a significant role in driving the speciation of neotropical fish species [18]. This extensive swampy system covered the western Amazon during most of the Miocene (around 10 to 20 mya) [15] and experienced marine incursions from the Caribbean, which likely influenced the adaptations of species from estuarine to freshwater conditions The Pebas mega-wetlands This supports the hypothesis that the colonization of inland waters originated from the north and the Caribbean Sea, resulting in the mixed sister clade position of *Apionichthys* relative to *Hypoclinemus* in our biogeographic reconstruction (Fig. 4). The formation of the Andes and Cordillera de Mérida disrupted the connection between Lake Pebas and the Caribbean. Additionally, the emergence of the Vaupes Arc, associated with global cooling and lower sea levels, increased sediment discharge from the Andes. The formation of the Acre system also contributed to this process [55, 56]. This pattern of colonizing freshwater environments in the Amazon region has been observed in other aquatic species, such as Amazonian pufferfish [57] and potamotrygonin stingrays [4].

**Fig 4.**
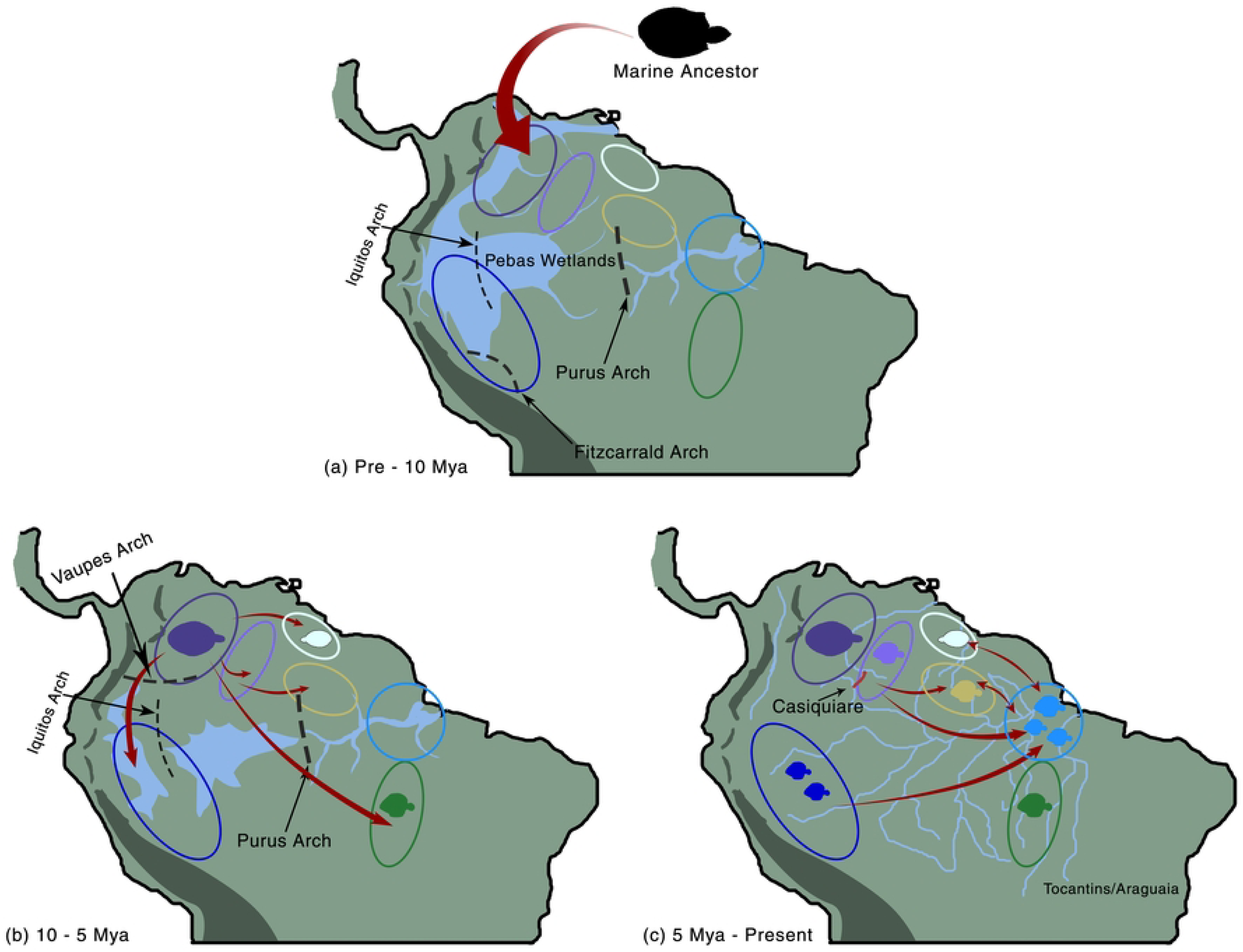
Graphical summary of overall changes in distribution of *Hypoclinemus* lineages over the time based on BBM analysis of RASP 4 proposed by our study. Red Arrows show the direction of colonization. Colors of ellipses correspond to the same ecoregions shown in Fig. 3. (a) Early/middle Miocene (Pre ∼10 mya); (b) Late Miocene/Pliocene (∼10-5 mya); and (c) Pliocene (5 mya – Present) distribution.

Before 10 million years ago (mya), the region now recognized as the lower Amazon was geographically separated from the upper Amazon by the Purus Arch. This geographical barrier created distinct eastern and western (Pebas) faunas [4, 58, 55] (Fig. 4a). The rise of the Vaupes Arch, approximately 10 mya (Fig. 4b), led to a substantial influx of sediments from the sub-Andean foreland and eventually caused the rupture of the Purus Arch [59].

The subsequent formation of the modern transcontinental Amazon during the mid-Pliocene, around 5.11 mya (Fig. 4c), allowed uninterrupted sediment flow from the Andes to the Atlantic [56]. This connection between the western and eastern parts of the Amazon basin facilitated the mixing of lineages in the lower Amazon. This aggregation was favored by the river capture system [60], which enabled the dispersal of species from different origins, including the Orinoco (within lineage 3), the Upper Amazon (from lineage 4 to lineage 6), and the Guyana Shield (corresponding to the center of origin of *Hypoclinemus* lineage 7). These populations remained separated from each other due to the uplift of the Mérida Andes [16, 55].

River rearrangements play a crucial role in isolating and promoting secondary contact among lowland Amazonian birds, which can lead to reticulation events resulting in species or populations that do not align with their geographic distribution [61]. In the case of *Hypoclinemus*, the apparent lack of significant morphological variation across its wide range, as observed in the lineages proposed here, could be attributed to low-frequency but continuous downstream dispersal. The recent divergence between lineages IV, V, and VI can be traced back to a shared ancestor across this extensive range. Since these lineages consist of populations from various regions, local adaptations may not become fixed due to genomic admixture. While *Hypoclinemus* does exhibit minor variations in color patterns (Figure 5), these differences do not appear to be strongly linked to geography or phylogenetic relationships.

**Fig 5.**
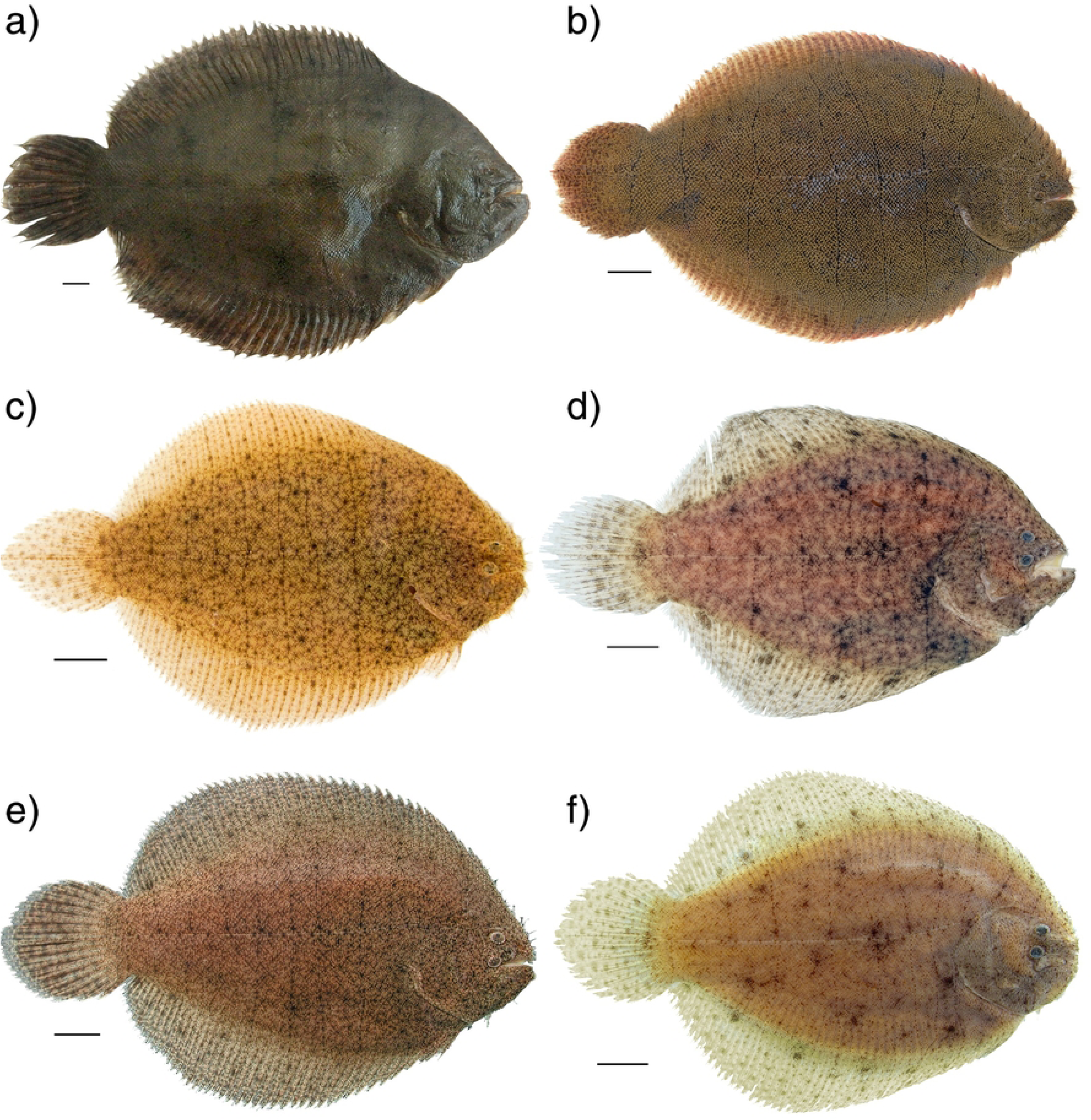
*Hypoclinemus mentalis* from throughout its distribution. A-individual from Brazil: Portel, Marajó Island (Lineage 5); B-Live specimen (IAvH-P 17110), Colombia: Meta: río Guayabero (Guaviare-Orinoco Dr.), 2°17’32.4” N, 73°52’31.6” W (Lineage 1); C-Live individual from lower rio Xingu, Brazil (undetermined between Lineages 1 or 5); D-Preserved specimen (ANSP 179511. 78.3 mm SL, tissue T2062), Guyana: Rupununi River (Essequibo Dr.) at Massara’s Landing, 3°53’41” N, 59°17’37” W (Lineage 5); E- Live individual from vicinity of Iquitos, Peru (Lineage 4); F-Preserved specimen (ANSP 163857, 80.5 mm SL), Venezuela: Amazonas: caño (Orinoco Dr.) downstream from Quiratare, 2°59’N, 66°4’W (Lineage 2). Photos by A. Guimarães (A), M. Sabaj (B, D-F) and L. Sousa (C). scale bar = 1 cm for A, D, F; unavailable for B, C, E.

The emergence of the Fitzcarrald Arch had a significant impact on the compartmentalization of the western Amazonas region, creating a separation between the Amazon River and the Madeira River [62]. This geological event is possibly associated with the speciation of lineage I (Xingu), as revealed in the present study. Over the past six million years, there have been periodic fluctuations in sea levels, particularly during the Pleistocene, when sea levels reached up to 120 meters at the end of the Pliocene [63, 64]. The Eastern Amazon was strongly affected by sea level cycles during the Pleistocene [4, 65, 66]. These sea level fluctuations may have facilitated intermittent connections between the ecoregions occupied by *Hypoclinemus* lineage VII, including Amazonas Estuary and Coastal Drainages, Amazonas Guiana Shield, Essequibo, and Xingu. These connections could explain the extensive distribution of this lineage. The transition period between Pliocene and Pleistocene led to the transcontinental formation of the Amazon River, which was affected by wide climatic variations with glacial and interglacial periods [67]. This period was marked by rapid speciation of the remaining lineages recovered in the present study, as isolation between the Xingu and Tocantins River basins.

Given all this, we proposed that the dynamics of Pebas wetlands during the Miocene plays a significant role for the dispersion of all cryptic lineages found in our study, where it’s created several connections between drainages, allowing fish faunal exchanges possible [56], together with the process of extension and contraction of headwater boundaries stablish connections between different river basins [11, 68]. After this broad dispersal process, the Pleistocene period promoted the speciation of *Hypoclinemus* species complex through allopatric speciation. The highlands acted as a refugee during glaciation period followed by colonization of lowlands, along with all actors relating here (e.g: marine incursions, uplift of paleoarches, historical connections promoting cross-drainage dispersal). All this evidence suggests a significant role of Pleistocene as a force that promotes speciation in neotropical fish species [4, 18, 58, 66, 69, 70), where *Hypoclinemus* is the more recent case.

## Conclusions

Our analysis reveals that *Hypoclinemus* is a monophyletic genus comprising at least five cryptic lineages. *Hypoclinemus* and *Apionichthys* form a predominantly freshwater clade. The speciation of *Hypoclinemus* appears to be influenced by the glacial cycles of the Miocene and Pleistocene, which suggests that freshwater taxa emerged during periods of marine incursions into the Pebas wetlands. The early diversification of *Hypoclinemus* lineages appears to be linked to major drainage regions, including the Orinoco, upper Amazon, and Guiana Shield/Lower Amazon, with subsequent downstream dispersal from the upper to lower Amazon and localized dispersal in the Guiana Shield/Lower Amazon region. The phylogeography of *Hypoclinemus* lineages indicates that dispersal has been a key factor in overcoming local genetic differentiation and likely contributes to the morphological homogeneity observed across the wide range of *Hypoclinemus*.

## Acknowledgments

This work has been supported by the National Council for Scientific and Technological Development (CNPq) through scholarships to AESR through orientation by JBLS as part of the Post-graduate Program in Aquatic Ecology and Fisheries (PPGEAP) and DJFS through orientation by JSR as part of the Post-graduate Program in Environmental Biology (PPGBA). We are deeply thankful for all lab support from Instituto Tecnológico Vale-Desenvolvimento Sustentável (ITV-DS). Fieldwork by MHS supported by the All-Catfish Species Inventory (NSF DEB-0315963) and iXingu Project (NSF DEB– 1257813). This work was also supported in part by the projects BRC16/19 (Biodiversity Research Consortium Brazil Norway Project 4183), SAMBA (RCN – INTPART Project number: 322457) and Centro de Triagem de Invertebrados (ITV-DS, Project Number: 4390). Additionally, the authors would like to thank Jamily Lorena de Lima and Luiz Filipe Brito de Oliveira for their help in the molecular procedures of the present study.

## Author Contributions

### Conceptualization

Alan Erik Souza Rodrigues, João Bráullio de Luna Sales.

### Data curation

Alan Erik Souza Rodrigues, João Bráullio de Luna Sales.

### Formal analysis

Alan Erik Souza Rodrigues.

### Funding acquisition

João Bráullio de Luna Sales, Jonathan Stuart Ready, Mark Sabaj.

### Methodology

Alan Erik Souza Rodrigues, João Bráullio de Luna Sales, Jonathan Stuart Ready.

### Sampling

Alan Erik Souza Rodrigues, Kamila de fatima Silva, Lucas Silva, Marcelo C. Andrade, Derlan Silva.

### Writing – original draft

Alan Erik Souza Rodrigues.

### Writing – review & editing

All authors.

---

**S1 Table.**
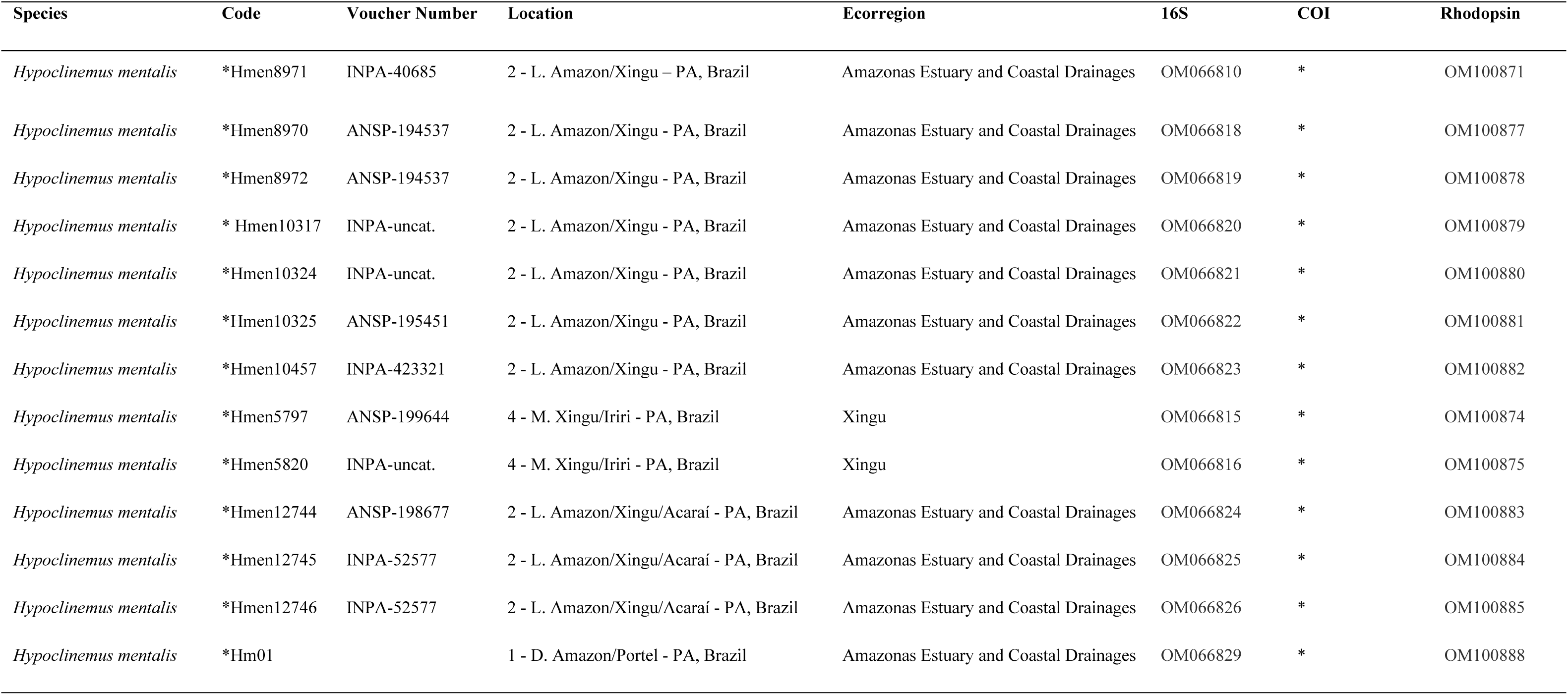
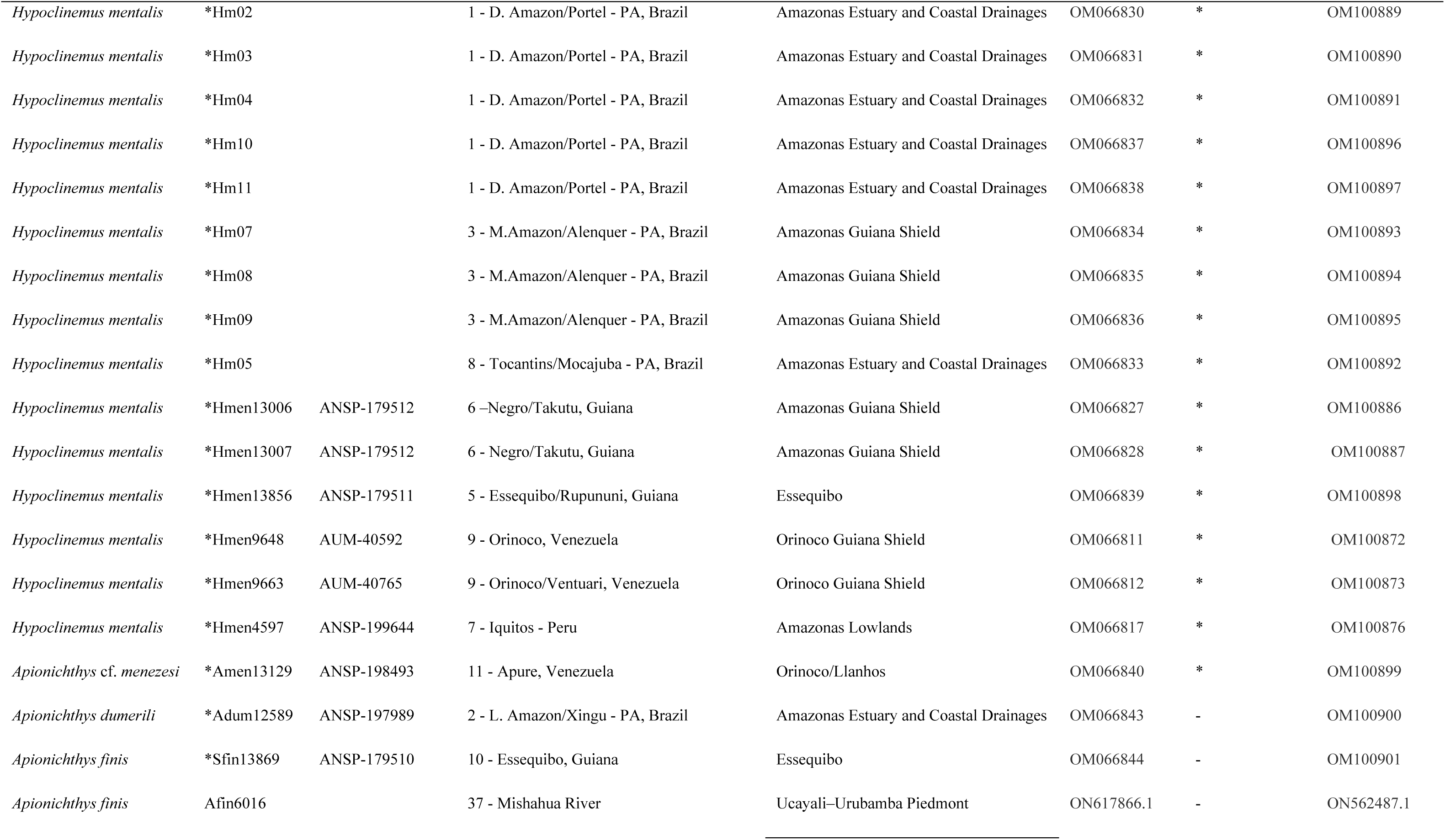
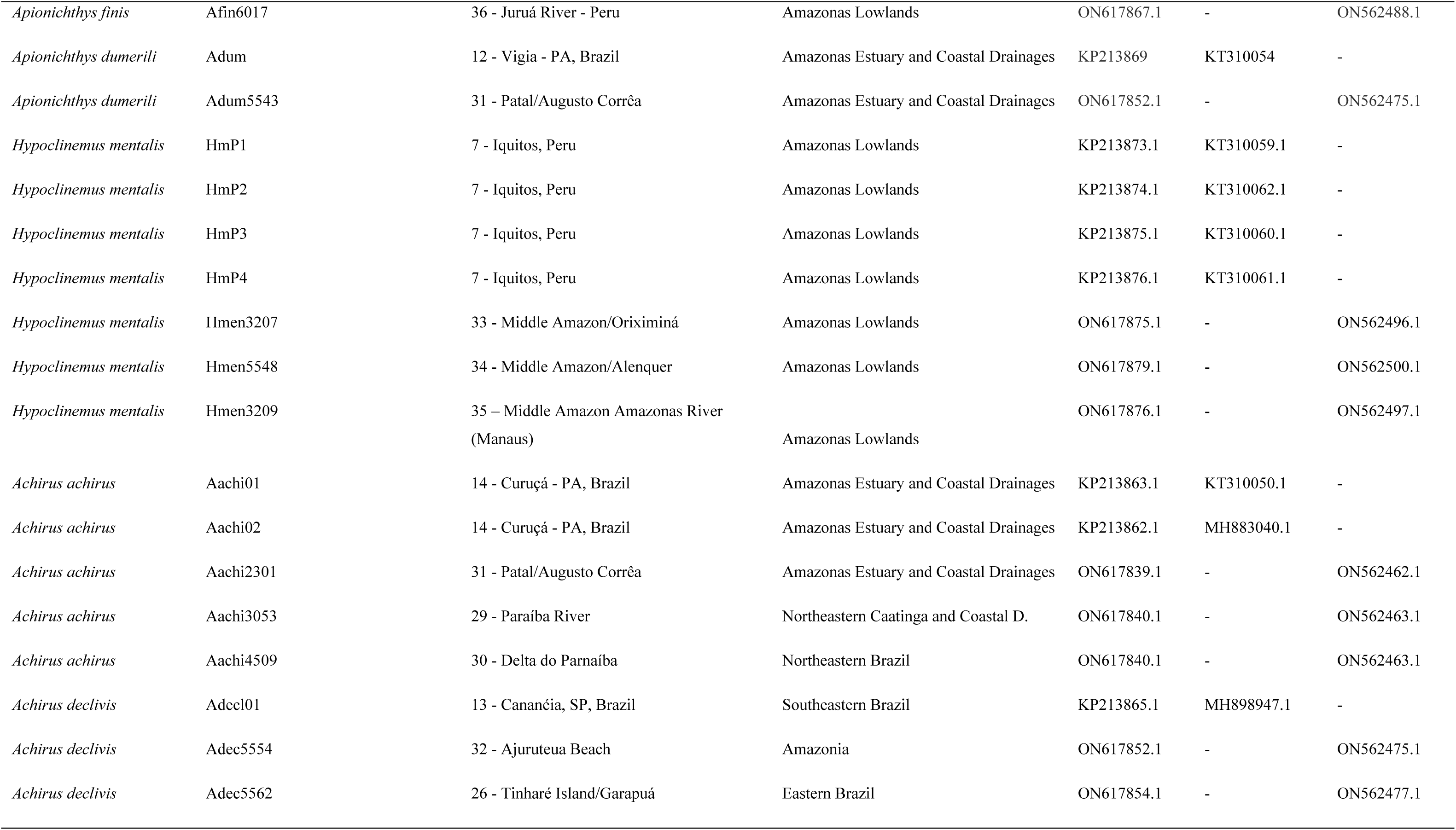
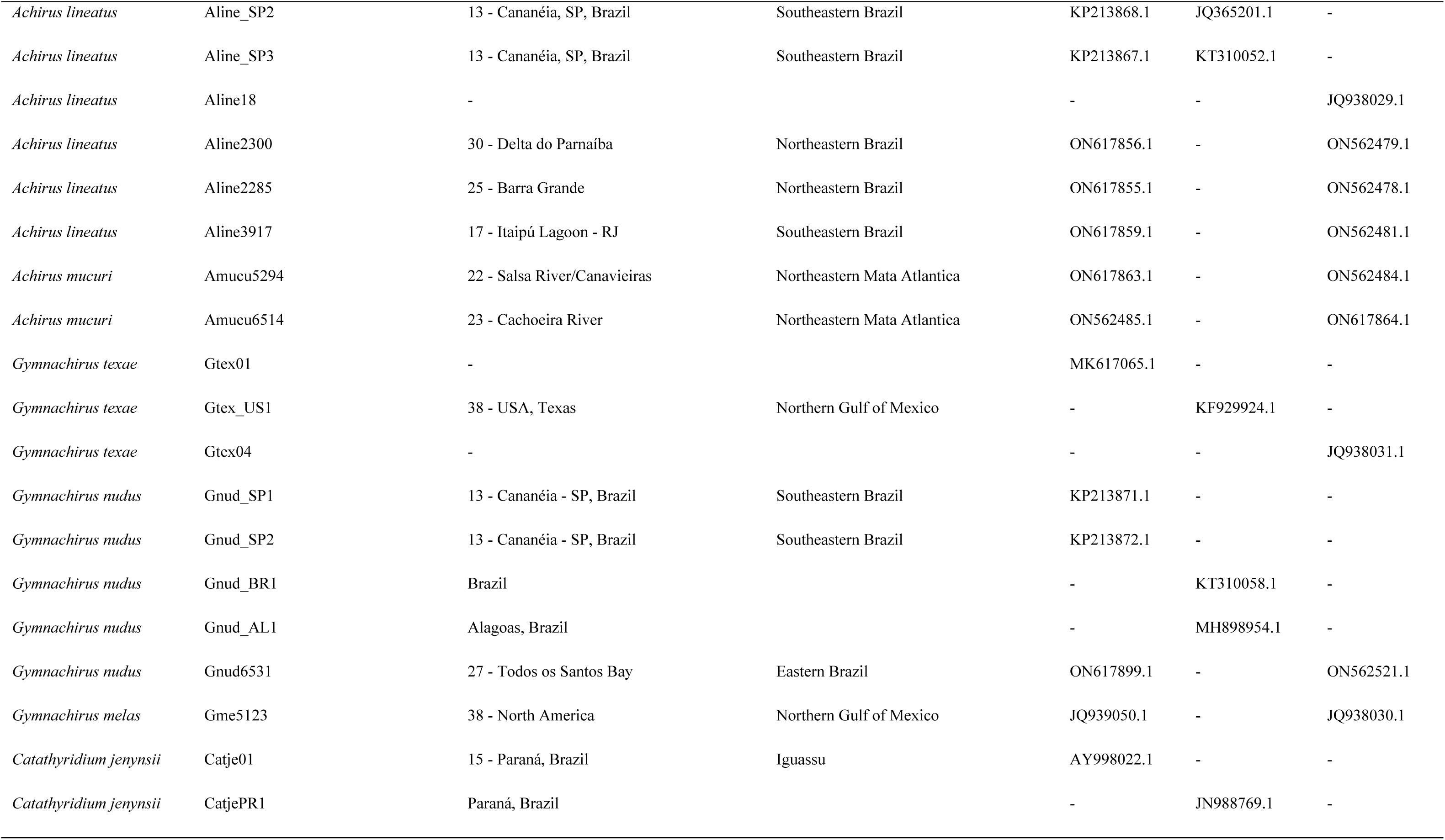
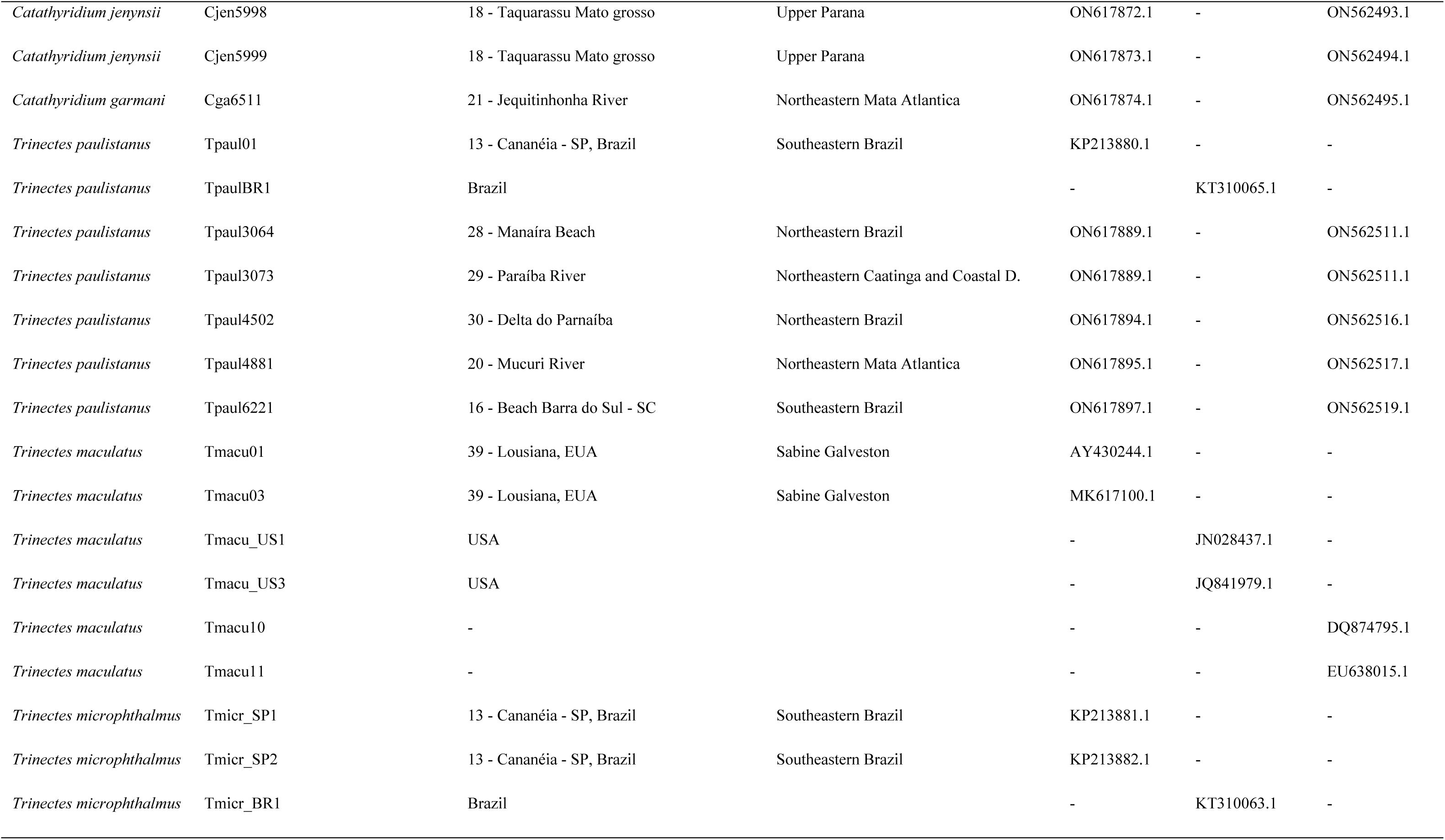
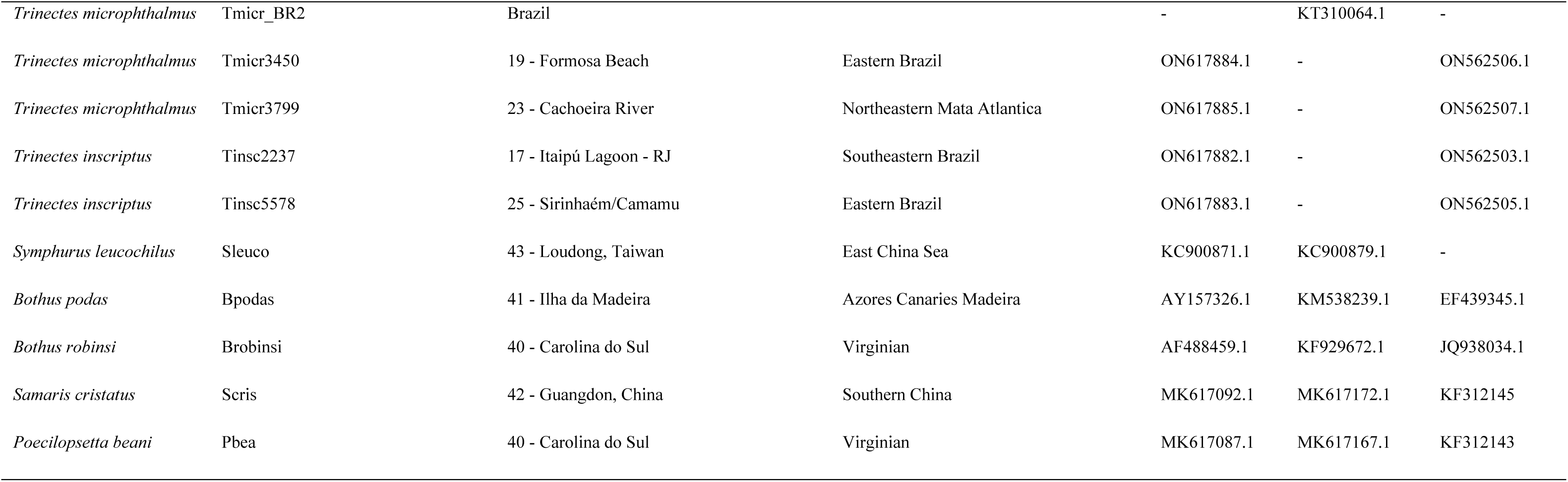
Species name, sample code, location, ecoregion and amplified regions. The numbers represent the respective locations in Fig. 1. Samples sequenced in the present study are indicated with an asterisk (*) with their respective markers. The sequence samples from Genbank are shown according to each gene and representative of the family Achiridae.

**S1 Fig**. **Bayesian inference (Bi) trees of *cox1* (A) and rrLn+*RHO* (B) of *H. mentalis* cryptic lineages recovered in the present study.** Only support values above 0.95 for Bi are shown and for maximum likelihood, only supports above 75% are indicated. Grey filled circles indicate *a posteriori* probabilities ≥ 0.8 (80%). Colors represent ecoregion associations inferred in RASP 4.

